# Integration of chemosensing and carbon catabolite repression impacts fungal enzyme regulation and plant associations

**DOI:** 10.1101/2021.05.06.442915

**Authors:** Wolfgang Hinterdobler, Guofen Li, David Turrà, Miriam Schalamun, Stefanie Kindel, Ursula Sauer, Sabrina Beier, Aroa Rodriguez Iglesias, Stéphane Compant, Stefania Vitale, Antonio Di Pietro, Monika Schmoll

## Abstract

Fungal metabolism and enzyme production are regulated by nutrient availability and by interactions with the living environment. We investigated the mechanisms underpinning adaptation of the biotechnological fungus *Trichoderma reesei* to decaying plant biomass versus living plants. We found that concentration-gated response to glucose, the main molecule sensed from dead plant biomass, is mediated by a conserved signaling pathway downstream of G protein-coupled receptors (GPCRs), while the carbon catabolite repressor CRE1 is critical for glucose concentration gating. The GPCRs CSG1 and CSG2 are further required for root colonization and formation of appressorium like structures on plant surfaces. Acceleration of sexual development in the presence of plant roots and their interactions with fruiting bodies indicates preferential association with plants. Our results reveal a complex sensing network governing resource distribution, enzyme production and fungal development that explains previously observed phenomena in fermentations and opens new perspectives for industrial strain improvement and agriculture.

## Introduction

Sensing of the environment is crucial for all living organisms – be it in their natural habitat or in an industrial setting. Fungi are efficient degraders of plant biomass, and some can also act as root symbionts of living plants by sharing critical nutrients and extending the plant’s acquisition network (1–3). The ability to distinguish dead plant material that needs to be enzymatically degraded from a living plant that serves as a mutualistic association partner, is essential to activate the appropriate metabolic and developmental responses during the interaction.

Fungi of the genus *Trichoderma* are ideal models for transfer of natural phenomena to industrial application by studying complex interspecies relationships, as they can act both as saprophytic biomass degraders and plant symbionts (4, 5). *Trichoderma reesei* (6, 7) is mostly known as an efficient degrader of cellulosic plant material in nature (8). However, this species was also shown to protect plants against a soil borne oomycete pathogen (9), suggesting that it can undergo beneficial associations with plant roots. Moreover, *T. reesei* is the only member of the genus in which sexual development has been achieved under laboratory conditions (10), which is significantly regulated by pheromone sensing and external cues (11) and of considerable biotechnological importance for strain improvement (11, 12).

Fungi can utilize insoluble plant biomass as a source of nutrients by secreting an array of cell wall degrading enzymes (CWDEs) (13). Regulation of CWDE expression in *T. reesei* has been studied in detail (4, 14). The disaccharide sophorose is thought to act as a natural inducer of cellulolytic enzymes (15, 16). However, in the presence of easily metabolized carbon sources such as glucose, CWDE synthesis is inhibited through the carbon catabolite repressor CRE1 (17, 18). Furthermore, regulation of cellulase genes in *T. reesei* requires conserved components of the heterotrimeric G protein/cAMP pathway, including the G protein alpha subunits GNA1 and GNA3, the G protein beta and gamma subunits GNB1 and GNG1, as well as the phosducin like protein PhLP1 (19–23). Recently, two class XIII G protein-coupled receptors (GPCRs), CSG1 and CSG2, were shown to be required for cellulase expression, and CSG1 was also implicated in posttranscriptional regulation of cellulase production (24).

Fungal hyphae have the capacity to chemotropically sense and follow a variety of directional cues, allowing them to successfully locate nutrient sources, mating partners or host organisms (25). Although fungal chemotropism has been known for more than a hundred years (26), the underlying mechanisms and cell signaling pathways remain poorly understood (25, 26). Recently, hyphae of the soil-inhabiting plant pathogen *Fusarium oxysporum* were found to grow towards a variety of carbon and nitrogen sources, as well as towards host plant signals. The chemotropic response to plants is triggered by class III peroxidases released by the roots and requires the fungal peptide pheromone receptor Ste2, as well as a mitogen-activated protein kinase (MAPK) pathway and the NADPH oxidase (NOX) complex (27, 28). While this groundbreaking discovery provided a first glimpse into the complexity of fungal chemosensing, little is known so far on chemotropic signaling in other fungi or on signal integration.

Here we report the identification of a chemotropic signaling network in *T. reesei*, which is able to precisely sense glucose concentrations and to accordingly modulate signaling outputs. We further establish that this mechanism allows the fungus to discriminate between dead plant material and living plants and to activate the appropriate metabolic and developmental responses, i.e. enzymatic degradation of biomass for nutrient supply versus root colonization and sexual development.

## Results

### *T. reesei* re-orients its hyphal growth towards gradients of nutrient sources

Chemotropic growth towards plant exudates has been reported in *F. oxysporum* (27) and *Trichoderma harzianum* (29). Here we used a previously described plate chemotropism assay to measure chemotropic sensing in the reference strain QM6a of *T. reesei* (Figure 1a). Because QM6a does not readily germinate in water agar, 0.0025 % w/v peptone was added to the agar. This concentration was optimized to ensure coordinated conidial germination above 60 % without interfering with chemotropism. As previously reported for *F. oxysporum* (27), *T. reesei* exhibited a significant chemotropic response to 1 % (w/v) glucose and 5 % (w/v) sodium glutamate (Figure 1b). These results confirm that the optimized assay conditions are appropriate for measuring the chemotropic response of *T. reesei* to different carbon and nitrogen sources.

**Figure 1.**
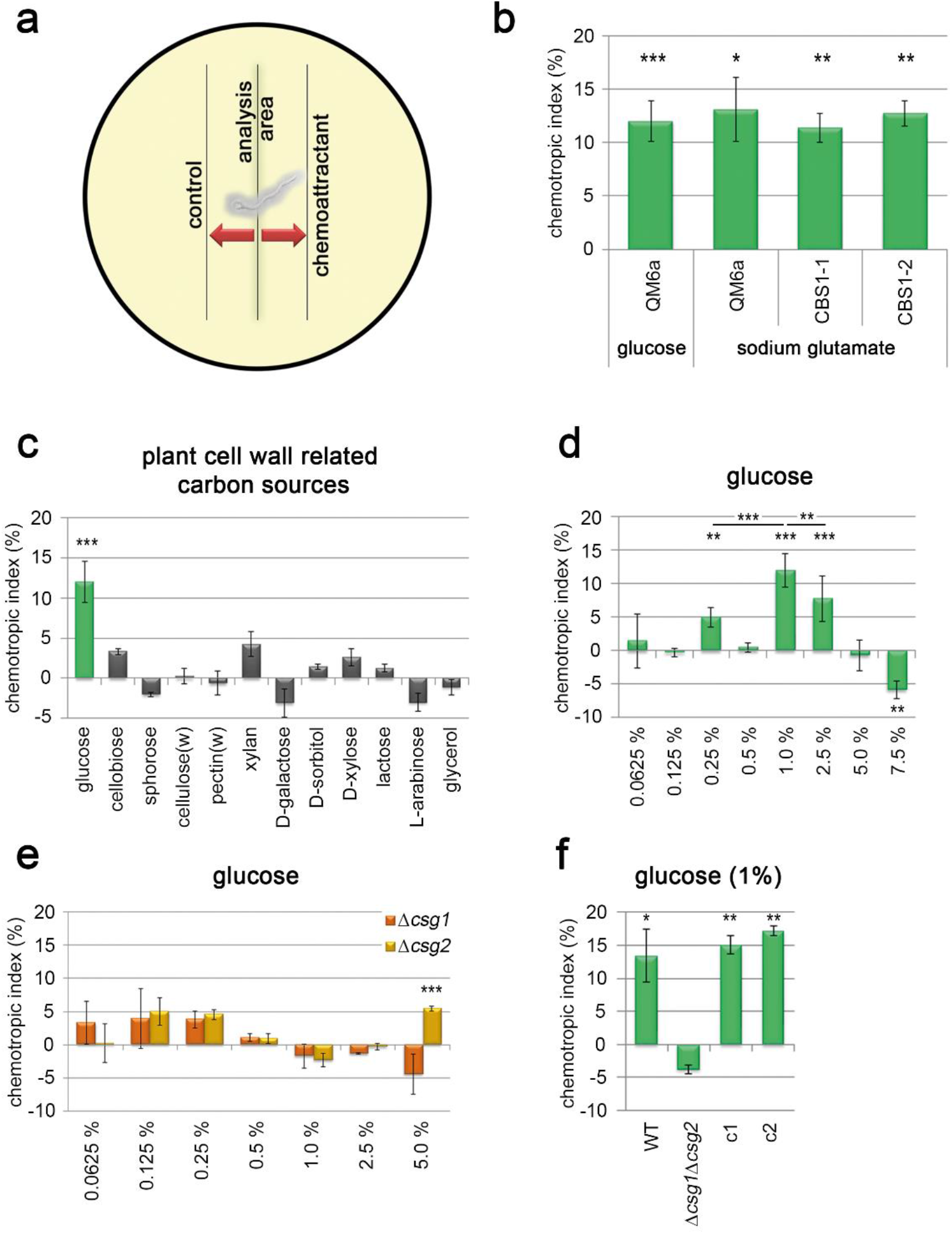
*T. reesei* responds chemotropically to nutrients. (**a**) Schematic representation of the experimental setup for analysis of chemotropic responses. (**b**) Chemotropic response of different *T. reesei* wild type strains to1 % glucose or 5 % sodium glutamate. (**c**) Chemotropic response of wild type strain QM6a to different carbon sources related to degradation of plant biomass. Soluble carbon sources were used at 55 mM corresponding to 1 % glucose, cellulose and pectin were used at 1 % and washed (“w“) after autoclaving prior to application. (**d, e**) Chemotropic response of QM6a (**d**) or the G protein-coupled receptor mutants Δ*csg1* and Δ*csg2* (**e**) to different glucose concentrations. (**f**) Chemotropic response of a Δ*csg1*Δ*csg2* double mutant and two progeny from the crossings of QM6aΔ*csg1* and QM6aΔ*csg2* with FF1, in which these deletions are restored (c1 and c2). Error bars show standard deviations from at least two biological replicates. Asterisks mark statistical significance of chemotropism in comparison to the control (absence of a chemoattractant). Statistical significance between tested strains or concentrations is indicated by asterisks over black bars. * p-value < 0.1, ** p-value < 0.05 and *** p-value < 0.01.

### Glucose is the main chemoattractant released from plant cell walls

CWDE expression in *T. reesei* is tightly regulated to trigger production of the appropriate enzyme cocktail for degradation of the different cell wall materials (15). We therefore asked whether this fungus can chemotropically sense building blocks of the plant cell wall. None of the high molecular weight compounds cellulose, xylan and pectin elicited a significant chemotropic response, most likely due to the insoluble nature of these polymers (Figure 1c). In line with this idea, cellulose and pectin did induce a significant chemotropic response after autoclaving (Figure S1a), and this response was abolished when the soluble compounds released by autoclaving were washed away prior to the chemotropism assay (see below). Hyphal chemotropism of *T. reesei* is thus elicited by soluble degradation products of cellulose and pectin, as previously reported in *F. oxysporum* (27).

We next measured the chemotropic response of QM6a to different glucose concentrations ranging from limiting levels (0.0625 % w/v) to very high concentrations (5 %) generating considerable osmotic pressure. An optimum chemotropic response was detected at 1 % (55 mM) glucose, with a gradual decrease at higher or lower concentrations (Figure 1d). The bell-shaped dose–response curve is reminiscent of that found in *F. oxysporum* (27) for peptide pheromones (30, 31). Testing of additional *T. reesei* strains confirmed a robust chemotropic response to glucose with a peak at concentrations between 0.5 and 1 % (Figures S1b and S1c). We conclude that glucose chemosensing in *T. reesei* is subject to gating, allowing for an optimum chemotropic response in a limited range (gate) of concentrations.

We next tested additional low molecular weight carbon sources released upon plant cell wall degradation or involved in enzyme regulation, including glycerol, cellobiose or lactose, the transglycosylation product sophorose, as well as to D-xylose, D-galactose or L-arabinose, all of which act as inducers or repressors of CWDEs (16, 32). However, none of these compounds elicited a significant chemotropic response when applied at 55 mM, which provided the optimum response for glucose (Figure 1c). Based on these results, we conclude that glucose is the main soluble compound released from degradable plant material eliciting chemotropic hyphal growth in *T. reesei*. Nevertheless, these other plant cell wall derived compounds may still be sensed by the fungus and cause physiological responses other than chemotropism.

### Glucose chemosensing requires the GPCRs CSG1/CSG2 and functions via the heterotrimeric G protein-cAMP-PKA pathway

We next asked how chemotropic sensing of glucose is mediated in *T. reesei*. Previous studies suggested that the two class XIII GPCRs CSG1 and CSG2, which are involved in regulation of cellulase gene expression, could act as cellulose or glucose-sensors (24). However, the precise function of these GPCRs in glucose signaling has not been addressed. Here we found that the chemotropic response to all glucose concentrations tested was largely abolished in the Δ*csg1* and Δ*csg2* single and double mutants (Figure 1e, f). By contrast, mutants in two other GPCR genes, Δ4508 and Δ80125, were unaffected in their response, thus confirming the specificity of CSG1 and CSG2 in glucose chemosensing (Figure S1d) and that the *hph* selection marker does not interfere with chemotropic responses. Similar to the wild type strain, the Δ*csg1* and Δ*csg2* mutants failed to show a significant response to the other low molecular weight carbon sources tested (Figure S2). Collectively, these findings suggest that CSG1 and CSG2 have essential and non-redundant roles in chemotropic sensing of glucose. This is in line with previous results showing that loss of CSG1 or CSG2 has differential impacts on cellulase regulation (24).

Heterotrimeric G protein subunits (19–21) as well as adenylate cyclase ACY1 and protein kinase A (22) were previously shown to function in cellulase regulation (Figure 2a). Here we found that the chemotropic response to 1 % glucose was abolished in *T. reesei* deletion mutants lacking either the G alpha subunits GNA1, GNA2, or GNA3, the G beta subunit GNB1, the G gamma subunit GNG1, the phosducin PhLP1, the catalytic subunit of protein kinase A, PKAc1, or the adenylate cyclase ACY1 (Figure 2 a, b). Interestingly, constitutive activation of GNA1 and GNA3 (19, 20) , but not of GNA2, also abolished the chemotropic response to glucose (Figure 2c). We conclude that the heterotrimeric G protein-cAMP pathway plays a key role in concentration dependent glucose chemosensing of *T. reesei* and that fine-tuned activation levels of the G alpha subunits GNA1 and GNA3 are required for this response.

**Figure 2.**
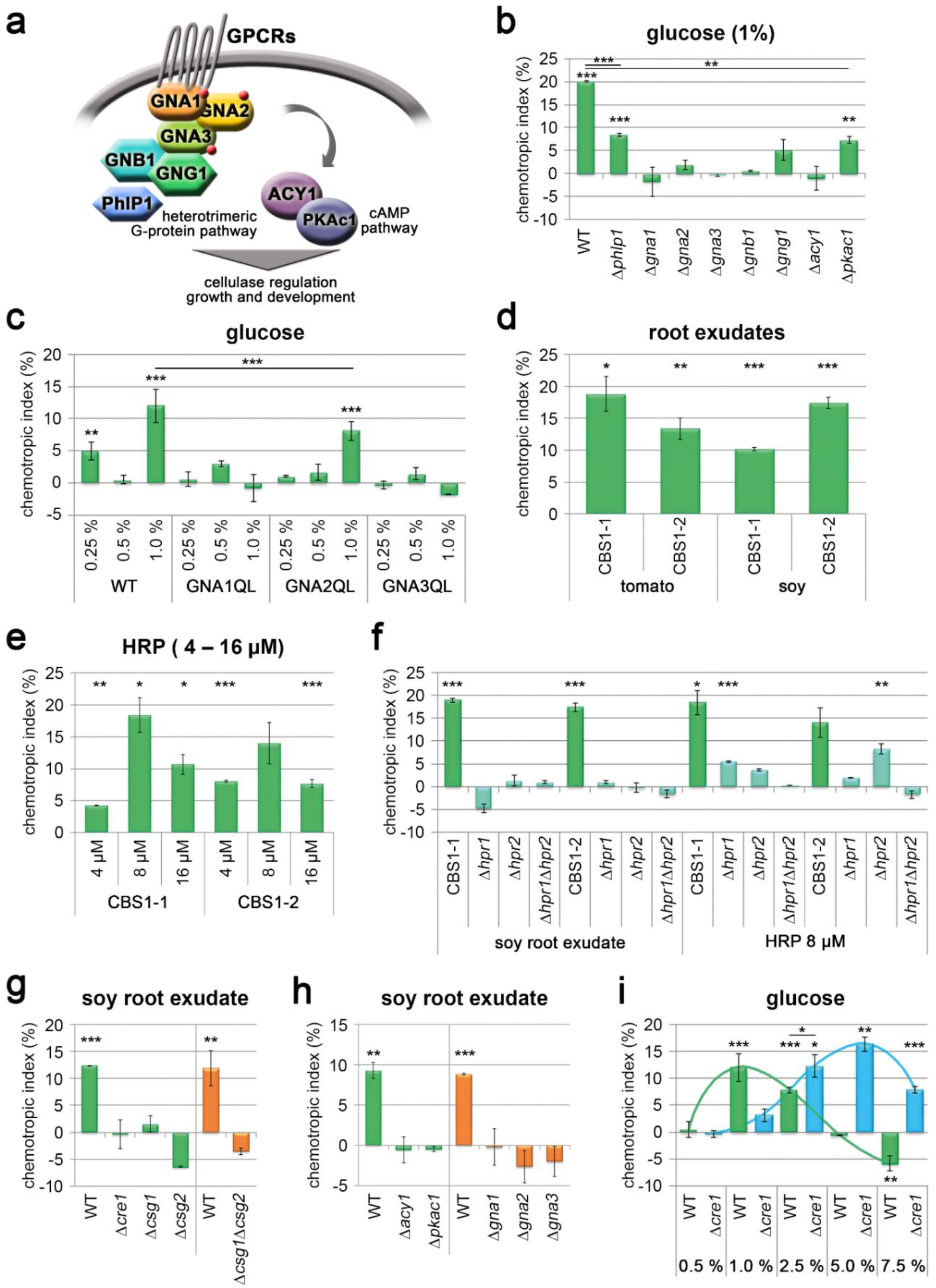
Glucose chemosensing requires the GPCRs CSG1/CSG2, heterotrimeric G proteins and the cAMP/PKA pathway. (**a**) Schematic representation of the heterotrimeric G protein-PKA/cAMP pathway in *T. reesei*. (**b**) Chemotropic response to 1 % glucose of deletion mutants lacking the phosducin PHLP1, G alpha (GNA1/2/3), beta (GNB1) or gamma (GNG1) subunits, the adenylate cyclase ACY1 or the catalytic subunit 1 of protein kinase A (PKAc1). (**c**) Chemotropic response to different glucose concentrations of strains carrying dominant activating alleles of the G protein alpha subunits GNA1, 2 or 3. (**d**) Chemotropic response to root exudates of tomato or soybean plants, of the fully fertile strains CBS 999.97 MAT1-1 (CBS1-1) and CBS999.97 MAT1-2 (CBS1-2). (**e**) Chemotropic response to different concentrations of horse radish peroxidase (HRP) of CBS1-1 and CBS1-2 . (**f**) Chemotropic response to soy root exudate or 8μ M HRP, of CBS1-1 or CBS1-2 or the single and double pheromone receptor deletion mutants derived from these strains. (**g**) Chemotropic response to soy root exudate of deletion mutants lacking *cre1*, *csg1* and/or *csg2*. (**h**) Chemotropic response to soy root exudate of deletion mutants lacking the indicated G alpha subunits, adenylate cyclase (ACY1) or the catalytic subunit 1 of protein kinase A (PKAc1). (**i**) Chemotropic response to different glucose concentrations of the wild type and the Δ*cre1* mutant. Error bars show standard deviations from at least two biological replicates. Asterisks mark statistical significance of chemotropism in comparison to the control (absence of a chemoattractant). Statistical significance between measurements is indicated by asterisks over black bars. * p-value < 0.1, ** p-value < 0.05 and *** p-value < 0.01.

### CSG1/CSG2 and the G-protein-cAMP-PKA pathway are also required for chemosensing of plant roots

Although *T. reesei* is mostly known as an efficient degrader of dead plant material (8), this fungus has been suggested to undergo beneficial associations with plant roots (9). We therefore asked whether *T. reesei* is able to chemotropically sense the presence of the living plant. In the plant pathogen *F. oxysporum*, chemotropic growth towards roots is triggered by secreted class III plant peroxidases and requires the GPCR Ste2, the cognate receptors of peptide sex pheromone alpha (27, 30). Here we found that similar to *F. oxysporum*, *T. reesei* exhibited a significant chemotropic response to root exudates from soybean or tomato plants (Figure 2d), as well as a concentration dependent response to commercial horse radish peroxidase (HRP) with a peak at 8 μM (Figure 2e). Both responses required the GPCR orthologues of Ste2 and Ste3, which are named HPR1 and HPR2 in *T. reesei* (Figure 2f). Importantly, HPR1 and HPR2 were also required for chemotropism of *T. reesei* towards peptide sex pheromones a and alpha, respectively (supplementary note 1, Figure S3), mimicking the results from *F. oxysporum*. Thus, the dual role of the pheromone receptors in chemosensing of mating pheromones and plant peroxidases is conserved between the two fungal species, reinforcing the functional link between sexual development and plant recognition in fungi.

We next asked whether, in addition to their role in glucose chemosensing, the GPCRs CSG1/CSG2 and the heterotrimeric G protein-cAMP-PKA pathway are also required for chemosensing of living plant roots. In contrast to the wild type strain, the Δ*csg1* and Δ*csg2* single and double mutants lacked a chemotropic response to root exudates (Figure 2g). Likewise, mutants in GNA1, GNA2, GNA3, PKAc1 or ACY1 were impaired in directed growth towards the chemoattractants released from plant roots (Figure 2h). Taken together, these results indicate that chemosensing of plant signals by *T. reesei* depends on the sex pheromone receptors as well as on the GPCRs CSG1 and CSG2, which act upstream of the G protein/cAMP pathway.

### The carbon catabolite repressor CRE1 determines chemotropic sensitivity to glucose and root exudates

Due to the importance of glucose as a signaling molecule released from plant biomass, we next investigated the role of carbon catabolite repression (CCR), a conserved regulatory mechanism that prevents unnecessary production of fungal CWDEs in the presence of readily utilizable carbon sources such as glucose (18). Similar to other fungi, CCR in *T. reesei* is mediated by the conserved carbon catabolite repressor CRE1 (17). Although the chemotropic response to glucose was still functional in a Δ*cre1* mutant, the optimum response concentration was dramatically shifted to 5 %, as compared to 1 % in the wild type strain (Figure 2i). At a higher concentration of 7.5 %, a decrease in the response was observed, suggesting that concentration dependence is still functional in the Δ*cre1* mutant. Most importantly, the Δ*cre1* mutant was also impaired in the response to plant root exudates (Figure 2g). Thus, CRE1-mediated carbon catabolite repression regulates the sensitivity of chemoperception and impacts gating of the chemotropic response both to glucose and to plant root exudates.

### Chemosensing via CSG1/CSG2 impacts hyphal morphology on cellulosic substrate as well as root colonization

In nature, nutrient sensing is crucial to induce adaptive responses to the substrate. For example, some *Trichoderma* spp. differentiate appressorium-like structures during mycoparasitic interaction with other fungi, thereby facilitating host adhesion and cell wall degradation by lytic enzymes (33, 34). Here we asked whether chemosensing through CSG1 and CSG2 impacts morphogenetic development of *T. reesei* in response to dead and living plant substrates. Scanning electron microscopy revealed the presence of characteristic appressorium-like attachment structures at the tips of the wild type hyphae growing on surface-sterilized maple leaves (Figure 3a). Similar structures were also observed during growth on cellophane membranes (Figure S4). However, no such structures were detected in the mutants lacking either CSG1 or CSG2 (Figure 3a, Figure S4).

**Figure 3.**
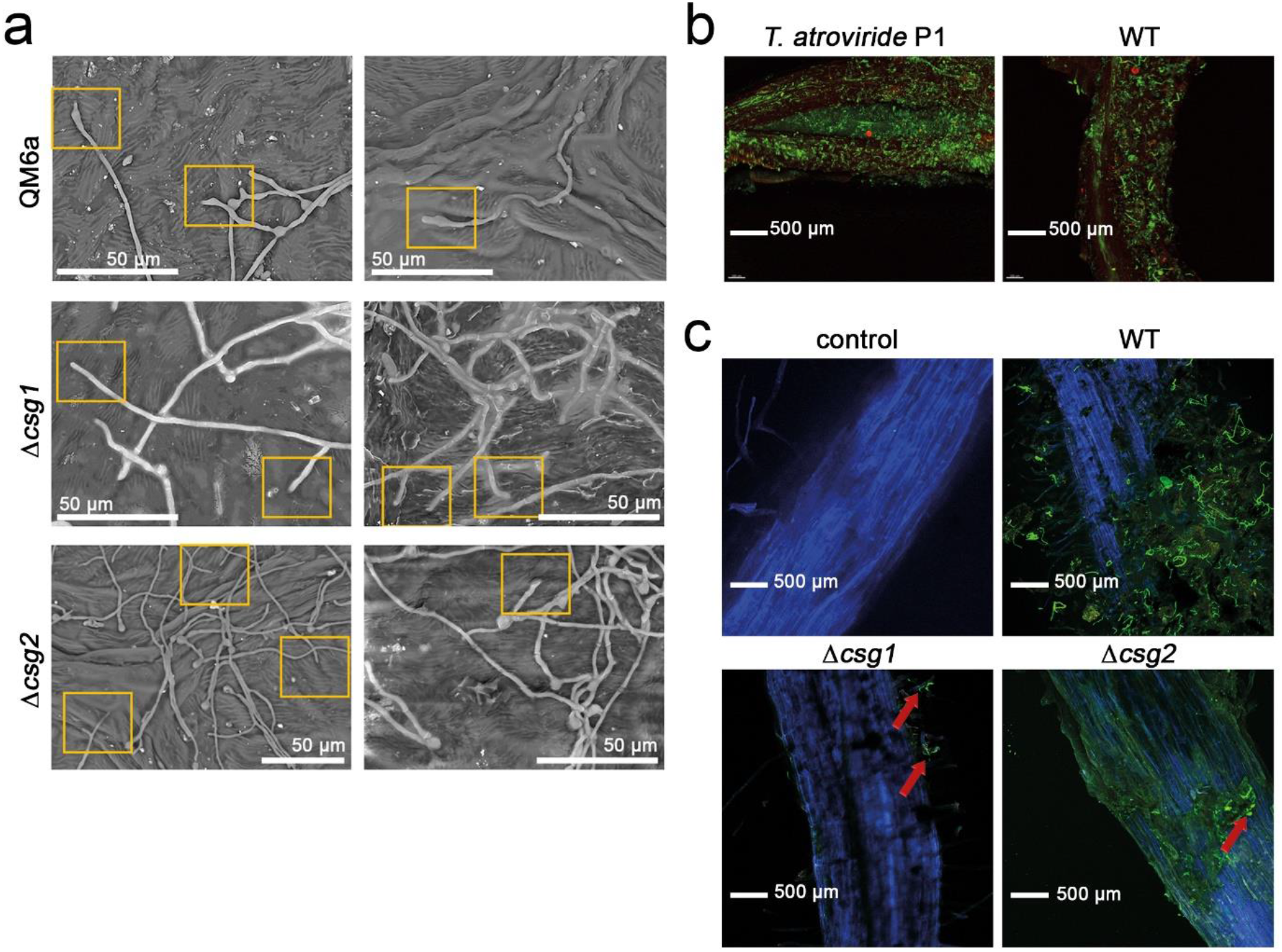
CSG1/CSG2 are required for developmental responses to plant surfaces. (**a**) Scanning electron microscopy analysis of interactions of QM6a and the receptor mutants Δ*csg1* and Δ*csg2* to a surface sterilized maple leaf. Yellow boxes indicate morphological alterations of hyphal tips in the wild type that are not observed in the deletion strains. (**b**) Confocal microscopy analysis of colonization of soybean roots by *T. atroviride* or *T. reesei* QM6a (WT). (**c**) Colonization of soybean roots by QM6a, Δ*csg1* or Δ*csg2*. An uninoculated root is shown as control.

We next tested the ability of the different strains to colonize living soybean roots. While the wild type strain of *T. reesei* colonized the roots nearly as efficiently as the well-known biocontrol strain *Trichoderma atroviride* P1 (Figure 3b), the Δ*csg1* or Δ*csg2* mutants largely failed in root colonization (Figure 3c). Collectively, these results suggest that chemosensing via CSG1/CSG2 not only mediates directed hyphal growth of *T. reesei* towards nutrient sources and plant roots but is also required for the differentiation of specialized adhesion structures on natural cellulosic substrates and for efficient colonization of living plant roots.

### Plant roots stimulate sexual development of *T. reesei*

Because nutrient availability is crucial for efficient sexual development (35), we wondered whether glucose sensing by CSG1/CSG2 impacts sexual development and/or chemical communication by secretion of secondary metabolites(36) prior to mating. Here we found that the mating behavior of the mutants lacking CSG1 and/or CSG2 was not significantly changed compared to wild type and that both showed altered profiles compared to wild-type with Δ*csg2* producing lower levels of sorbicillin derivatives (yellow pigments) (Figure S5a-h).

Our finding that chemotropic growth of *T. reesei* towards roots requires the Ste2 and Ste3 orthologs HPR1 and HPR2 confirms earlier results showing that chemotropic sensing of the host plant functions via a sex pheromone receptor (27, 30). We therefore asked whether plant signals impact sexual development of *T. reesei*. To this aim, mating experiments were performed in the absence or presence of soybean seedlings, using two different combinations of opposing mating types: strains CBS999.97 MAT1-1 × CBS999.97 MAT1-2 and FF1 × FF2, two female fertile derivatives of QM6a. In both mating assays, fruiting body formation was accelerated by one day in the presence of soybean seedlings (day 6 vs. day 7 in control conditions) (Figure S5g and Figure S6, 7). This effect was only observed in the physical presence of the plant, but not when plant root exudates were added to the medium. Moreover, accelerated fruiting body formation was not mimicked by the presence of a wooden toothpick or a glass rod, suggesting that it is not due to the presence of a physical structure on the plate (supplementary note 2 and Figures S8 and S9). The fruiting bodies produced in the presence of soybean seedlings were often larger in size and irregular in shape compared to those produced in the absence of the plant (Figure S6). Similar to the wild type, single mutants in pheromone receptors also showed accelerated fruiting body formation in the presence of soybean seedlings (Figure S5g). By contrast, as reported previously, fruiting body formation in a Δ*hpr1* Δ*hpr2* double receptor mutant was impaired and ascospore formation was abolished in the single receptor mutants of their cognate mating type (37). These defects were not rescued by the presence of the plant (Figure S7, red background). We conclude that interaction with the plant stimulates fruiting body formation in *T. reesei* but cannot substitute for the requirement of the pheromone-receptor interaction.

### *T. reesei* fruiting bodies physically interact with plant roots

To further explore the effect of the living plant on fungal mating, we conducted microscopic analysis of fruiting bodies forming on and around plant roots (Figure 4). We noted that these structures were firmly attached to the root surface and contained morphologically normal perithecia (Figure 4b–n). We further observed a thickening of the root around the contact zone with the fruiting body and could clearly discern an interaction zone, where hyphae extended towards the root without extensively penetrating it (Figure 4e, k, l). In cases where the roots grew across a fruiting body, deformation of both the root and the fruiting body was observed (Figure 4b, g–k, n), with the root remaining intact for at least 7 days after fruiting body formation. By contrast, roots which entered a fruiting body without exiting underwent progressive shrinking and degradation of the root tip inside the fruiting body, with fungal hyphae occupying the resulting cavity (Figure 4 d,e). Viability staining with methylene blue highlighted a region of the fruiting body mycelium surrounding the root canal, indicative of a response of the fungal mycelium to the plant (Figure 4e (4)). A striking response observed was the *de novo* formation of fruiting body tissue in the presence of the root (Figure 4b, g–i). Roots enclosed inside such a structure displayed signs of degradation, possibly as a result of plant or fungal defense reactions (Figure 4l–n). We next asked whether the absence of the fungal pheromone receptor would result in increased damage or growth reduction of the plant roots, but found no evidence for this (supplementary note 3, Figure S10). We conclude that fruiting bodies of *T. reesei* respond morphologically to presence of plant roots, which may act both as a carbon source as well as a stimulus for sexual development (see supplementary note 4).

**Figure 4.**
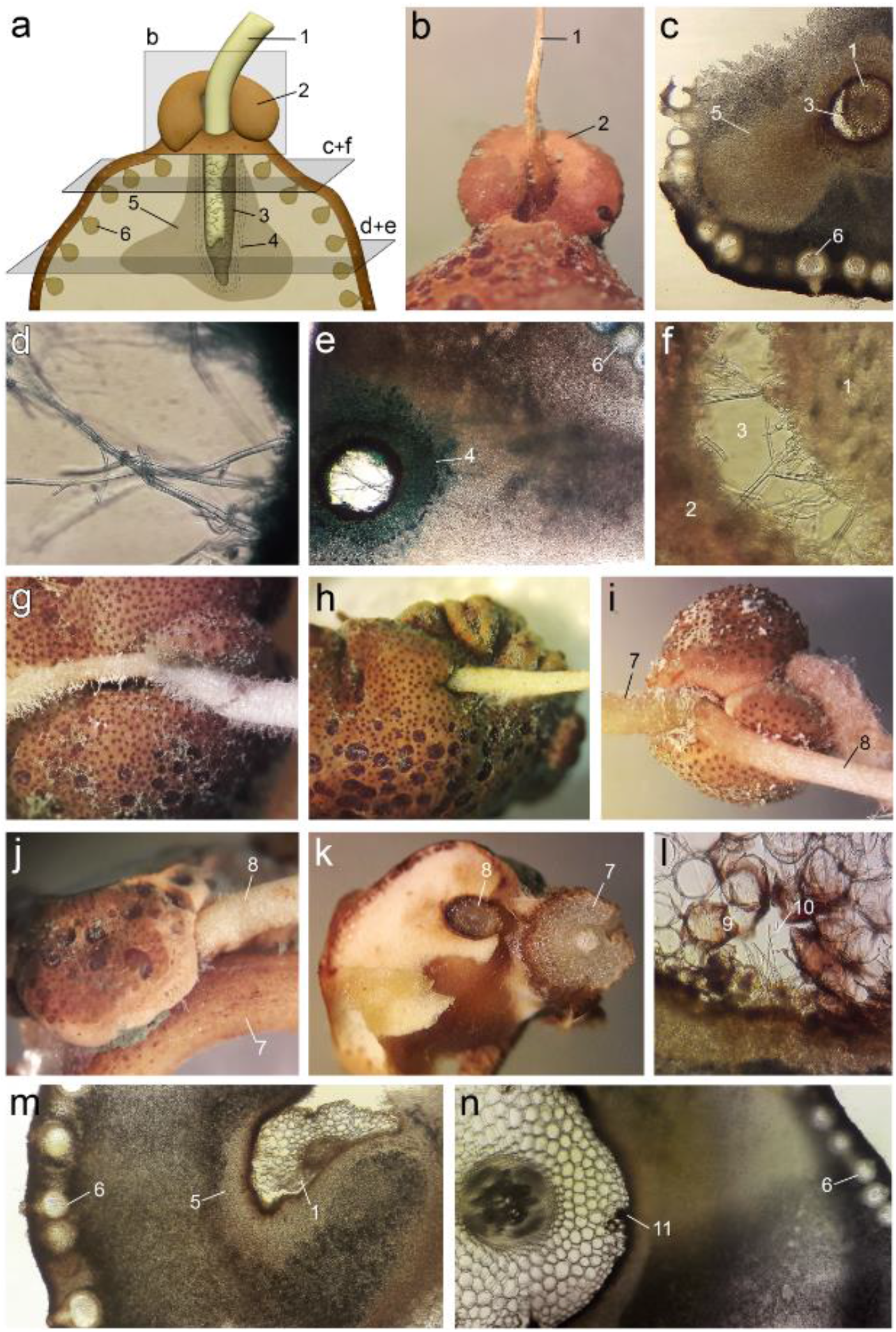
Fruiting bodies physically interact with soybean roots. (**a**) Schematic representation of the cross-sections through a fruiting body (FB) enclosing a soybean root [1]. Highlighted in grey are the macroscopic view of the entering zone (b), and two cross-sections (c) and (e) (200x) with the corresponding magnifications (d) and (f) (400x); Perithecia [6], zone in the FB showing a response to the soy root [5], methylene blue-stained pattern [4] and the cavity remaining after root tissue degradation [3]. (**b**) Example of a FB forming additional tissue [2] as reaction to the presence of the plant root [1]. (**g**–**i**) Macroscopic interaction between FB and roots; primary [7] and secondary roots [8].(**j** and **k**) FB grown on the branching area of primary [7] and secondary root [8]. (**l**) Hyphae entering the root tissue [10] with root cells reacting to the fungus [9] are shown. (**m** and **n**) Cross-sections through FBs growing around roots with an entering point of mycelium [11] into the root tissue. Photos (b, g–j, m and n) are composed of several merged pictures (focus stacking) for better depth of focus and in case of (m) and (n) for a broader overview of the whole cross-section.

## Discussion

Every single spot in nature is teeming with life, with diverse organisms competing or living in synergy. The natural habitat of *Trichoderma* are forests, where decaying plant material coexists with living plants (1). In this complex environment, the fungus must adapt its lifestyle to different necessities. Here we provide evidence showing that *T. reesei* integrates chemosensing of glucose monomers released from cellulosic plant biomass with signals from living plants, via the GPCRs CSG1/CSG2 and the conserved heterotrimeric G protein-cAMP-PKA signaling cascade, to efficiently adjust its responses to the surrounding nutritional and “social” landscape. To achieve this adaptation, our data indicate that *T. reesei* uses feedback cycles of CDWE release, sensing of degradation products such as glucose or sophorose, and subsequent adjustment of CDWE regulation (Figure 5). While the regulatory mechanisms controlling expression of CWDEs, sexual development and plant interactions in *Trichoderma* spp. have been studied in detail (4, 5, 11, 14), the biological relevance and the interconnections between the different pathways have remained elusive. Previous work indicated that CSG1 is crucial for posttranscriptional regulation of cellulase gene expression (24), but not for induction of transcription. We show that CSG1 senses a defined concentration of glucose, and that a specific signal level transmitted through the G-protein cascade is required for the chemotropic response. Consequently, sensing of this specific glucose concentration must be the signal required for initiation of enzyme production after transcriptional induction by sophorose and hence represents a second checkpoint. This explains why constitutive activation of G alpha subunits (19, 20), which, as shown here, are crucial for glucose signal transmission, does not result in inducer-independent cellulase production: without transcriptional induction (first checkpoint), glucose levels sensed by this pathway are not relevant. Thus we propose that, in agreement with previous work (24), chemosensing of glucose concentration represents a crucial checkpoint for efficient utilization of cellulosic substrates and establishment of a physical interaction with plant roots.

**Figure 5.**
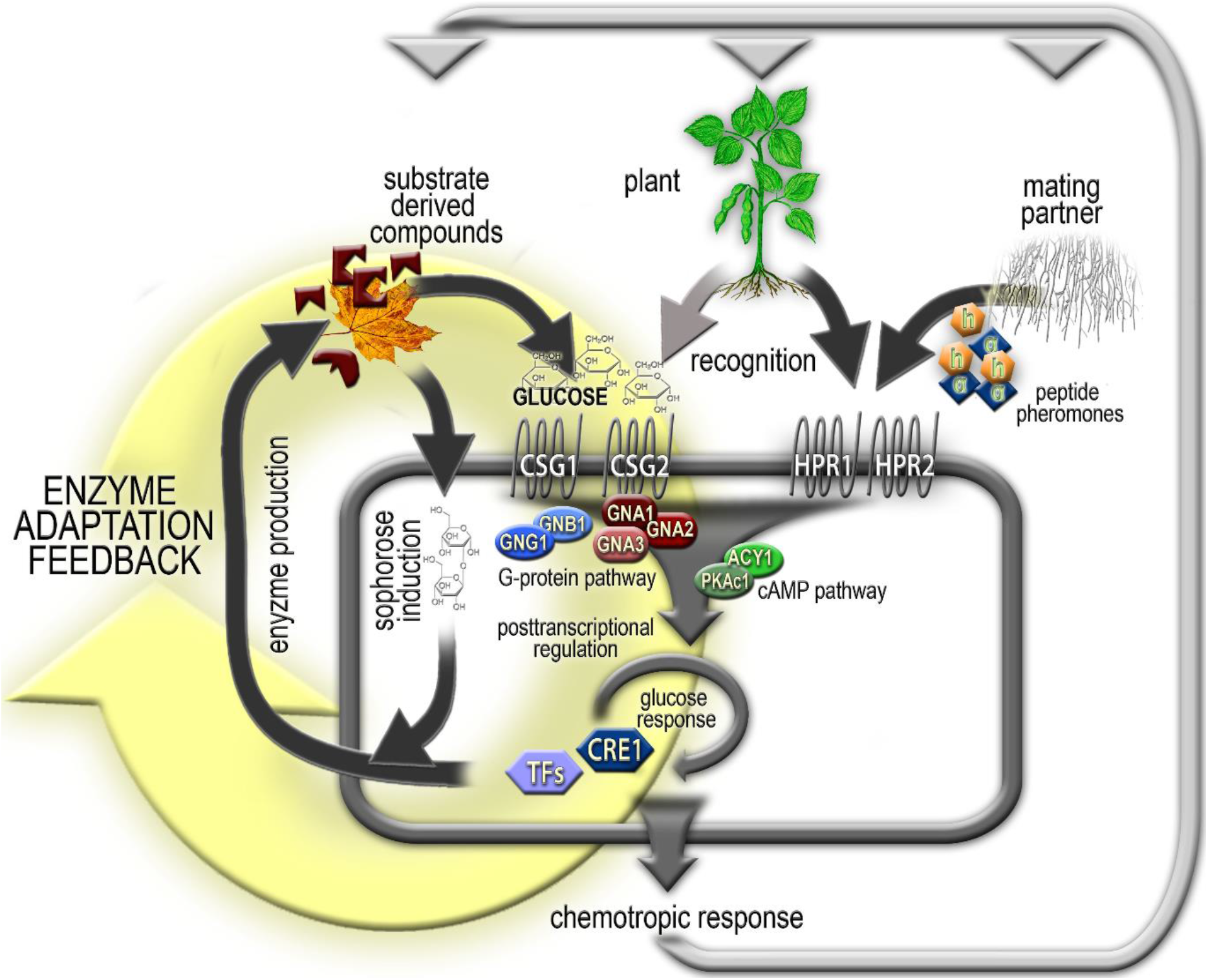
Integrative model of nutrient, plant and mating partner chemosensing in *T. reesei*. Transcription of genes encoding plant cell wall degrading enzymes (CWDEs) is triggered by the natural inducer sophorose. Moreover, CWDE biosynthesis beyond basal levels is initiated in response to defined levels of glucose liberated by enzymatic cleavage from plant biomass, requires the glucose sensor GPCRs CSG1/CSG2 and the heterotrimeric G protein-cAMP/PKA pathway and is negatively regulated by the carbon catabolite repressor CRE1 in response to high glucose levels. Proper nutrient sensing further results in morphogenetic responses and successful root colonization. Together these regulated pathways (enzyme secretion, glucose sensing, carbon catabolite repression and plant sensing) constitute an adaptation feedback loop which is used to distinguish between dead plant biomass (enzymatic degradation and glucose levels balanced) and living plants (enzymatic degradation not required for glucose liberation, no balance). The heterotrimeric G protein pathway integrates nutrient, plant and mating signals. Accordingly, transmission of nutrient signals by CSG1, CSG2 and CRE1 is required for proper plant recognition and sexual development is accelerated upon communication with the plant.

We further show that the carbon catabolite repressor CRE1 is important for concentration dependent glucose sensing. CRE1 transcript levels (38) and nuclear localization (39) were previously shown to be regulated by glucose concentrations. Our results are in line with previous findings showing that deletion of *cre1* alleviates the requirement for nutrient limitation to produce high enzyme levels in fermentors (40–42). Reminiscent of the “gating” found here for the chemotropic response to glucose, cAMP was shown to stimulate cellulase gene expression in *T. reesei* only at certain concentrations (23). Moreover, in yeasts glucose sensing and signaling was previously linked to the cAMP-PKA pathway (43, 44). Here we provide further support for the crucial role of the adenylate cyclase ACY1 and PKAc1, together with CRE1, in concentration dependent glucose signaling, which is likely to be crucial for improvements towards reliable upscaling of fermentations.

Degradation of dead plant material by fungal CWDEs yields increasing amounts of glucose, which are in balance with the level of CWDE gene expression. Our results suggest the presence of a negative feedback mechanism which is controlled via the fine-tuned sensing of glucose levels, in parallel to the detection of the inducer. This finding is biologically highly relevant, because living plant roots are known to secrete sugars (45) and exuded glucose could thus act as an inducing signal for root colonization by fungi (46, 47) in which transmission of nutrients to a fungal interaction partner is tightly regulated and requires contact (48, 49). Such a chemotropic signaling circuit should act separately from that controlling CWDE production to allow differentiation between dead litter and living plants acting as a “live nutrient source”. We propose that *T. reesei* distinguishes between live and dead plant material by sensing the balance of secreted CDWEs and the amount of glucose released by these enzymes. Combined with our findings that root signals are sensed via two different pathways - the pheromone receptors and the glucose-sending GPCRs CSG1/2, and that plant roots stimulate sexual development and interaction with fungal fruiting bodies, we propose that *T. reesei* applies a bipartite checkpoint mechanism involving glucose sensing via the heterotrimeric G protein pathway and the peroxidase signals detected by sex pheromone receptors. This mechanism is likely to enable *T. reesei* to optimize resource efficiency by tight regulation of CWDE biosynthesis versus plant association, which suggests that the latter is a preferable lifestyle for this fungus. This complex sensing mechanism has a high potential for understanding and improving fungal fermentations, strain improvement by crossing and exploration of fungal symbiosis or pathogenesis of plants.

## Methods

### Strains and cultivation conditions

For analysis of mating responses, recombinant strains in the fully fertile background of CBS999.97 in both mating types were used (10, 37). Assessment of chemotropic responses to nutrients were performed with wild type strains and mutants in the QM6a (6), QM9414 (50), TU-6 (51) or CBS999.97 strain backgrounds, depending on the mutants used, in order to be able to evaluate the results against the scientific background on regulation of plant cell wall degradation. Strains FF1 or FF2 (fully fertile derivative strains of QM6a; mating type MAT1-1 (FF1) or MAT1-2 (FF2)) were used for preparation of double mutants of selected genes (52). Strains Δ4508 and Δ80125 were constructed as described previously (53) and deletions were confirmed by PCR with primers binding within the deleted region. *Trichoderma atroviride* P1 (ATCC74508 (54)) was used as a control in the analysis of colonization of soybean roots. A list of the strains and genotypes used in this study is shown in Table S1.

For all chemotropic assays, strains were grown on malt extract agar (3 % w/v) at 28 °C until sporulation (4 days) prior to harvesting of conidia. For sexual development, compatible strains were cultivated on malt extract agar (2 % w/v) at room temperature in daylight (light-dark cycles) until fruiting body formation and subsequent ascospore discharge (55). For analysis of the relevance of soybean seedlings for sexual development, soybeans were surface sterilized, after 3 hours soaking in water, by brief immersion in 70 % ethanol, 2 minutes in 1.4 % sodium hypochlorite solution and two times washing with sterile water. Seeds were subsequently incubated on malt extract agar plates. Germinating soybeans were placed onto mating cultures prior to inoculation of fungal mating partners. All chemicals used were supplied by Roth (Karlsruhe, Germany) unless stated otherwise.

### Construction of Δcsg1 and Δcsg2 double mutants

The mutants Δ*csg1* and Δ*csg2* in the QM6a background (24) were crossed with strain FF1 and ascospores were harvested. Single spores were isolated and tested for the presence of the mutations by PCR. Mating types were analyzed as described previously (10). The resulting progeny for both mutants in female fertile background were crossed to obtain Δ*csg1*Δ*csg2* double mutants.

### Preparation of chemotropic agents and plant root exudates

For carbon sources, compounds were dissolved in purified water to the respective concentration and filter sterilized. Insoluble carbon sources were autoclaved and subsequently washed to remove degradation products released by autoclaving.

For preparation of plant root exudates, soybeans were surface sterilized (as described above) and planted in sterilized perlite (premium perlite 2–6, Gramoflor GmbH, Germany). Plants were grown until the second leaf stage and recovered from the perlite. After gentle washing with water, plant roots were carefully submerged in sterile water and kept for 2 days at room temperature. Root exudates were filter sterilized and stored at −80 °C.

### Analysis of chemotropic responses

Chemotropism plate assays were done essentially as described (27). Briefly, fresh conidia harvested from 4-day-old plates were resuspended in 1 mL spore solution (0.8 % (w/v) NaCl and 0.05 % (w/v) Tween 80). The suspension was filtered through glass wool, centrifuged at 8000 rpm for 2 minutes, the supernatant was discarded, and the pellet was resuspended in 1 mL of sterile purified water. The suspension was applied at a concentration of 10^8^ conidia per mL to assay plates containing water agar (0.5 % w/v; Roth, Karlsruhe, Germany No 5210.2). Different concentrations of peptone (from casein; Merck, Darmstadt, Germany; No 1.11931) were added to facilitate germination. The optimal concentration was determined to be 0.0025 % (w/v) peptone. For the uridine auxotropic strain TU-6, 10 μM uridine was added to the medium, when used as wild-type control. After application of the chemoattractant and the control solution, plates were incubated at 28 °C in darkness. After 13 hours of incubation, orientation of germ tubes was determined microscopically (VisiScope TL524P microscope; 200x magnification) and chemotropic indices were calculated as described earlier (27).

### Statistics

Statistic evaluation of results was performed using Student’s T-Test (two-sided) in R Studio (compare means, ggpubr v0.3.0). Background in variations of hyphal orientation of wild type strains in the absence of a chemoattractant was used as control for statistical evaluation (10 biological replicates).

### Analysis of patterns of secreted metabolites

Patterns of secreted metabolites were analyzed by excising agar slices close to the contact zone from plates with fungi grown under the conditions described. Three biological replicates of three plates pooled each were used for analysis as described previously by HPTLC (high performance thin layer chromatography) (56). Briefly, agar slices were extracted with chloroform and acetone, evaporated and resuspended in chloroform. The samples were spotted on TLC plates (HPTLC silica gel 60 F254s) using the CAMAG Automatic TLC sampler 4 (CAMAG, Muttenz, Switzerland). After separation, plates were analyzed at different wave lengths using the CAMAG visualizer (CAMAG).

### Analysis of colonization by *T. reesei*

Soybean seeds were surface sterilized and rinsed three times with PBS (phosphate buffered saline, pH7.2). Seeds were then put onto PDA plates containing *T. reesei* strains. Thereafter, the seeds were replaced into sterile magenta boxes containing twice autoclaved soil (1:1:1 perlite, sand, potting soil) and sterilized tap water (25 mL). After 10 days of incubation in the greenhouse, plants were harvested, immersed in PBS (15 mL) containing 50 μg/mL of wheat germ agglutinin (WGA)-AlexaFluor488 conjugate (Life Technologies, USA). Plant colonization was analyzed after 2 hours of incubation with (WGA)-Alexa Fluor488 at 37 °C and again rinsing three times with PBS.

Microscopy was performed using a confocal microscope (Olympus Fluoview FV1000 with multi-line laser FV5-LAMAR-2 and HeNe(G)laser FV10-LAHEG230-2, Japan) with objectives of 10x, 20x and 40x magnification. X, Y, Z pictures and scans were taken at 405, 488 and 549 nm with same settings and normal light. The software Imaris (Bitplane, Zürich, Switzerland) was used for visualization. ImageJ software (1.47v) was applied to merge pictures from different channels.

### Electron microscopy of substrate sensing

Leafs from *Acer pseudoplatanus* (sycamore maple) were used for analysis of hyphal morphology. Leafs were sterilized with 70 % (v/v) Ethanol and 10 % (v/v) DanKlorix solution (Henkel, Düsseldorf, Germany) washed three times with sterile purified water and dried at 65 °C over night. Electron microscopy was performed at a Hitachi Tabletop Microscope TM330 from a sample area of 5 × 5 mm using a cooling stage in the specimen chamber and freezing at −20 °C in the charge-up reduction mode to minimize distortions of images. At least three areas were investigated per sample in biological duplicates and at least one independent repetition was done with similar results.

## Author contributions

WH contributed to chemotropic analysis, performed microscopy and secondary metabolite analysis, participated in designing figures, performed statistical analysis of data and contributed to drafting and editing the manuscript. GL contributed to chemotropic analysis, growth assays and performed confocal microscopy under the supervision of SC. SC supervised work and designed pictures from confocal microscopy. SK and US performed electron microscopy. MiS and SB contributed to chemotropic analysis and SB further supported plant assays. ARI contributed construction of the strain with constitutive activation of GNA2. SV provided preliminary data and contributed to conception of the study. DT and ADP supervised part of the work of GL, participated in conception of the study, interpretation and editing of the final manuscript. MS conceived the study, supervised students, contributed to analysis of results, interpreted data, designed figures and wrote the final version of the manuscript.

## Acknowledgements

We want to thank Viktoria Fabsits for technical assistance and Stefan Böhmdorfer (University of Natural Resources and Life Sciences, Vienna) for the permission to use the HPTLC equipment for secondary metabolite analysis. Work of WH was supported by the NFB (NÖ Forschungs- und Bildungsges. m. b. H.; grant LSC16-004 to MS), work of GL, ARI, MiS and SB was supported by the Austrian Science Fund (FWF, grants P24350, P26935 and P31464 to MS). Work of SK was supported by the FEMtech program “Talents” of the Austrian Research Promotion agency (FFG). Work of DT was supported by grant BIO2016-78923-R from the Spanish Ministerio de Economía y Competitividad (MINECO) to ADP. We want to thank Markku Saloheimo (VTT) for providing the Δ*cre1* strain. Soybean samples from organic seed production were obtained from RWA Austria, Korneuburg, Austria, which is gratefully acknowledged.

## Competing interests

The authors declare no competing interests.

## Notes

### Competing Interest Statement

The authors have declared no competing interest.

