## Supplementary Material for "Integration of chemosensing and carbon catabolite repression impacts fungal enzyme regulation and plant associations"

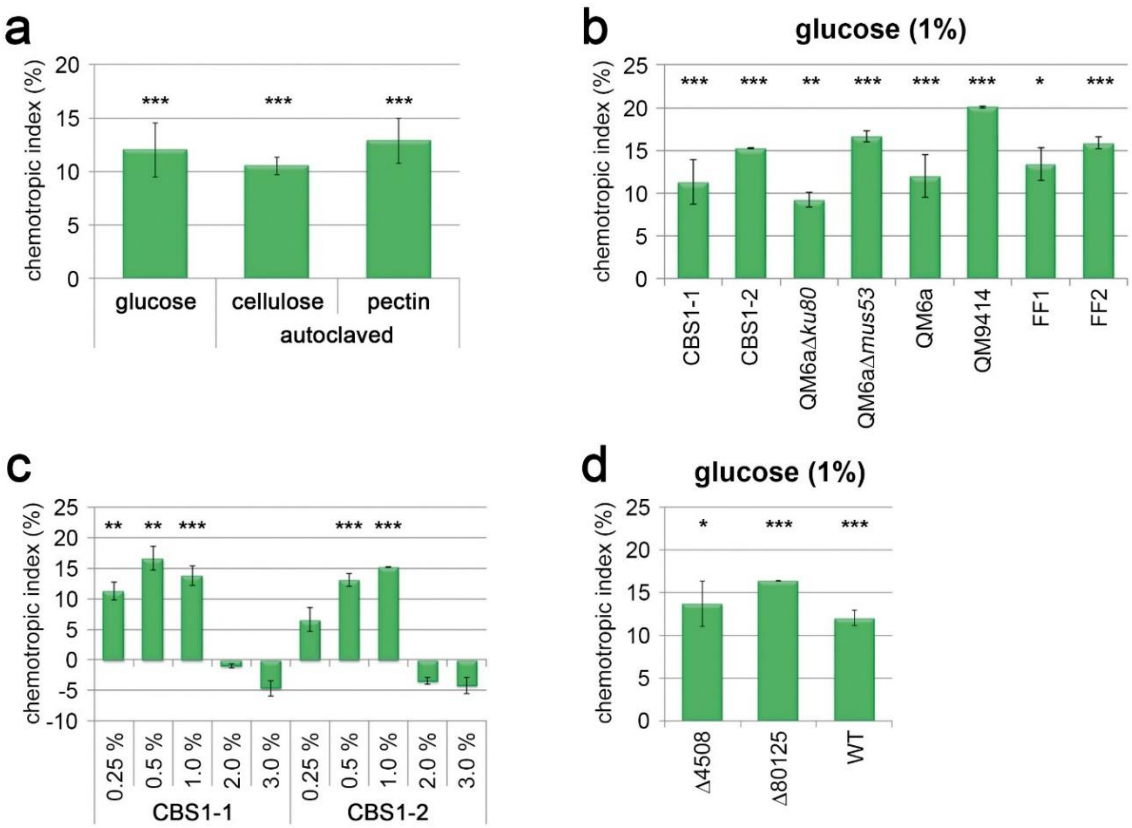


**Figure S1.** (**a**) Chemotropic response to autoclaved cellulose and pectin (without washing for removal of degradation products). (**b**) Chemotropic response of wild types and control strains to 1% (w/v) glucose. (**c**) Concentration dependence of chemotropic glucose sensing in CBS999.97 MAT1-1 and CBS99.97 MAT1-2. (**d**) Chemotropic response of strains harboring the *hph* selection marker to 1% (w/v) glucose. Here, we tested whether the presence of the hygromycin resistance gene in our deletion mutants affects the chemotropic response, and found that the GPCR mutant strains Δ4508 and Δ80125, all of which contain the *hph*-selection marker cassette, showed a similar response as the wild type strain. Error bars show standard deviations of at least two biological replicates. Error bars show standard deviations of at least two biological replicates. Asterisks mark statistical significance of chemotropism in comparison to background hyphal orientation of wild type strains in the absence of chemotropic agents. Statisitical significance between measurements is indicated by asterisks over black bars. * p-value < 0.1, ** p-value < 0.05 and *** p-value < 0.01.

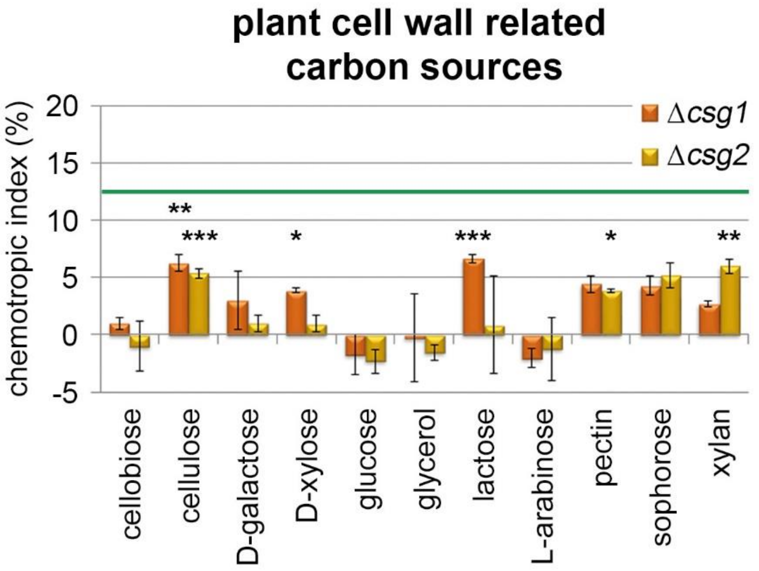


**Figure S2**. Chemotropic response of Δ*csg1* and Δ*csg2* to different carbon sources. The green line shows the level of chemotropic response of the wild type to 1% glucose. Error bars show standard deviations of at least two biological replicates. Error bars show standard deviations of at least two biological replicates. Asterisks mark statistical significance of chemotropism in comparison to background hyphal orientation of wild type strains in the absence of chemotropic agents. Statistical significance between measurements is indicated by asterisks over black bars. * p-value < 0.1, ** p-value < 0.05 and *** p-value < 0.01.

### Supplementary note 1

*T. reesei exhibits a chemotropic response to peptide mating pheromones*

In *F. oxysporum*, the pheromone receptor Ste2 is required for chemotropic sensing of peptide pheromone alpha, but also of plant roots (1). *T. reesei* has a recently discovered sexual cycle, which is triggered by peptide pheromones that signal via cognate GPCRs (2, 3). In this fungus, the function of the prenylated a-type pheromone is taken over by an unusual h-type pheromone whose mature, bioactive form is still unknown (4). We thus set out to test the role of peptide pheromones and their cognate receptors in the chemotropic response to a mating partner. A peptide encompassing part of the sequence of the a pheromone precursor (ERKRLIGCSVMTPKA) which previously showed bioactivity (5), triggered a concentration-dependent chemotropic response with an optimum concentration of 400 µM (Figure S4A). Importantly, this response was dependent on the presence of the cognate MAT1-1 pheromone receptor HPR1 (Figure S6A). However, since the synthetic peptide is not prenylated, as expected for natural a-type peptide pheromones, the underlying mechanism requires further study.
Unfortunately, the solubility of complete synthetic *T. reesei* alpha pheromone (4) (predicted sequence WCYRIGEPCW) was insufficient to induce significant chemotropy levels (Figure S6B). We therefore tested synthetic *F. oxysporum* alpha peptide pheromone (6) at its reported optimum concentration (1), and observed a significant response in type CBS MAT1-2, which was dependent on the presence of the Ste2 GPCR homologue HPR2 (Figure S4C and D). By contrast, with the *F. oxysporum* a-type peptide pheromone only a minor response below significance level was observed in both mating types. These results reveal a certain level of conservation between the pheromone sensing and response machineries of *Fusarium* and *Trichoderma*. Moreover, they confirm the functionality of the pheromone receptors HPR1 and HPR2 in pheromone sensing of *T. reesei* as well as a relevance of the HPP1 peptide ERKRLIGCSVMTPKA in the pheromone response.

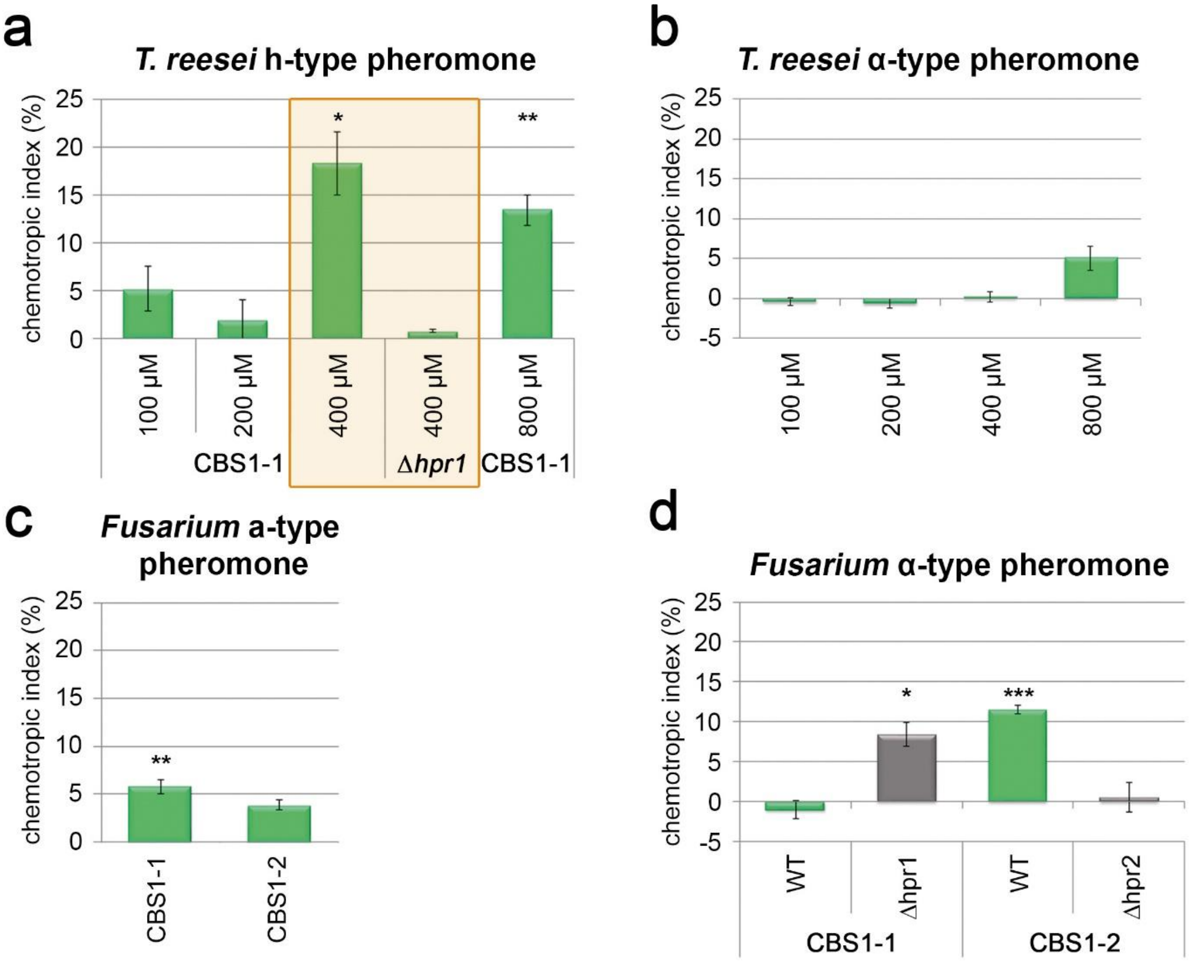


**Figure S3 Pheromone sensing by HPR1 and HPR2.** Chemotropic sensing was tested in the respective known cognate mating type(2) (h-type peptide pheromone assuming a-type function(4) with MAT1-1 and Δ*hpr1*) of *T. reesei*. For the *Fusarium* pheromones, both mating types were tested and pheromone receptor mutants were tested in the cognate mating types. (**a**) Chemotropic response of CBS999.97 MAT1-1 (CBS1-1) and a deletion mutant of the pheromone receptor gene *hpr1* in this strain background to different concentrations of a predicted peptide of the h-type peptide pheromone precursor HPP1 (4). The optimal concentration is boxed. (**b**) Chemotropic response of CBS999.97 MAT1-2 (CBS1-2) and to different concentrations of a predicted peptide of the α-type peptide pheromone precursor PPG1 (4). (**c**) Chemotropic response of CBS999.97 MAT1-1 (CBS1-1) and CBS999.97 MAT1-2 (CBS1-2) and the a-type peptide pheromone precursor of *F. oxysporum*. (**d**) Chemotropic response of CBS999.97 MAT1-1 (CBS1-1) and CBS999.97 MAT1-2 (CBS1-2) and to deletion mutants of the cognate pheromone receptor genes *hpr1* and *hpr2* in the respective strain background to the α-type peptide pheromone precursor of *F. oxysporum*. Error bars show standard deviations of at least two biological replicates. Asterisks mark statistical significance of chemotropism in comparison to background hyphal orientation of wild type strains in the absence of chemotropic agents. Statisitical significance between measurements is indicated by asterisks over black bars. * p-value < 0.1, ** p-value < 0.05 and *** p-value < 0.01.

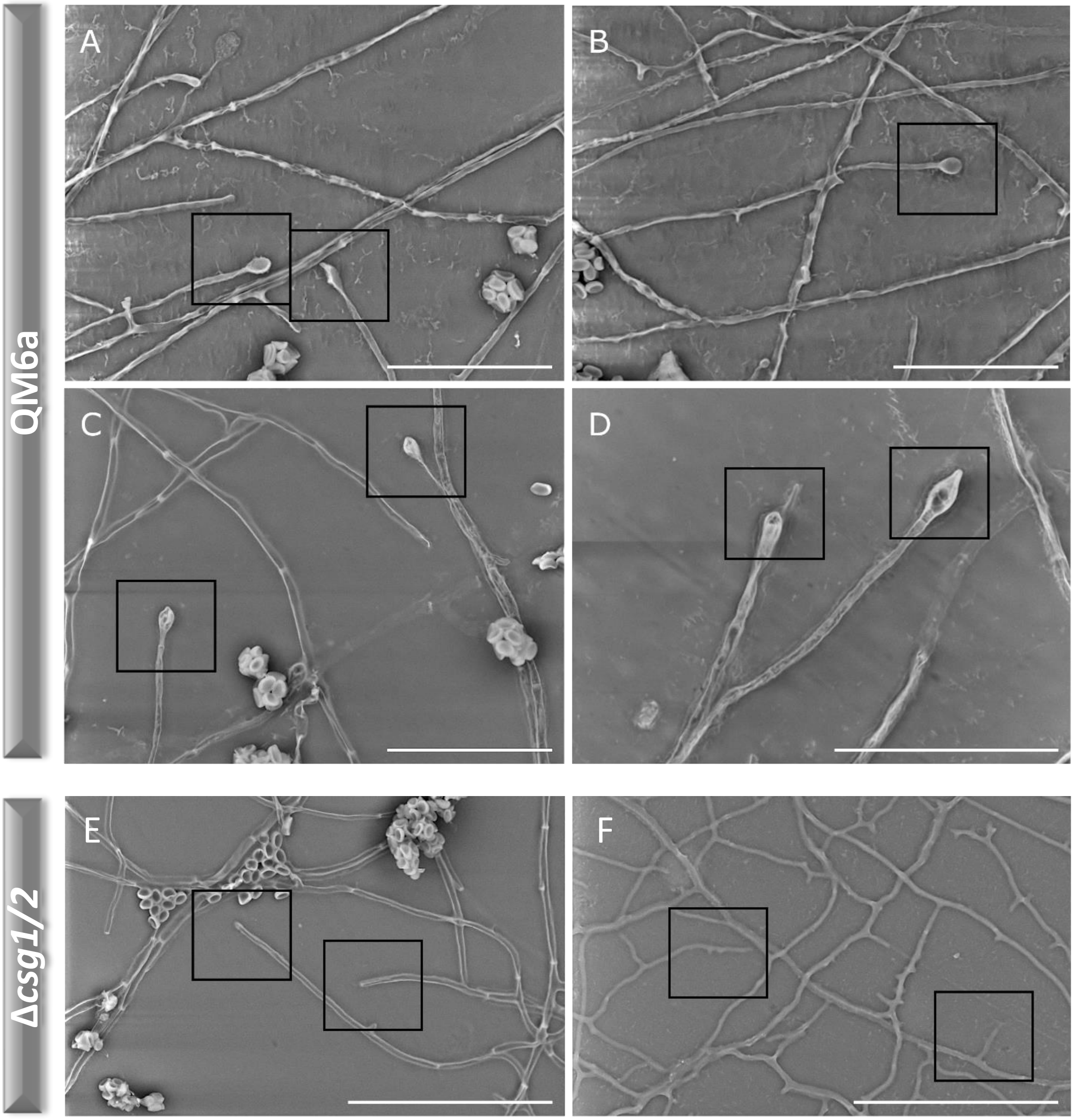


**Figure S4**. **Attachment structures formed on the surface of cellophane.** Scanning electron microscopy analysis of interactions of QM6a (a–d) and the G protein-coupled receptor mutants Δ*csg1* (e) and Δ*csg2* (f) to an agar surface overlaid with cellophane after 63 hours of growth. Boxes indicate morphological alterations of hyphal tips in the wild type that were not observed in the deletion strains. (d) moreover shows an attachment structure that appears to develop a hyphae reminiscent of infection hyphae. However, penetration assays remained negative.

(a–d) Scale bars = 30 µm, (e–f) Scale bars = 50 µm; Experiments were done in duplicates.

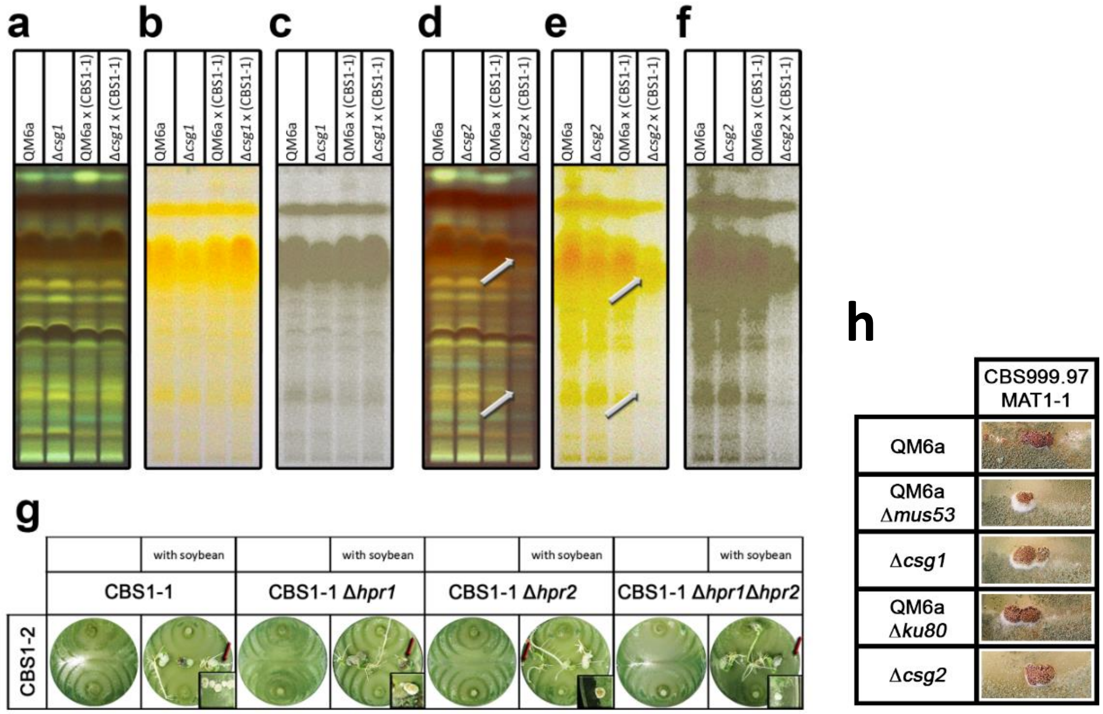


**Figure S5. Sexual development, plant and mating related responses.** (**a**–**f**) Secondary metabolite patterns of strains grown on malt extract agar (2 % w/v, light-dark cycles, 22 °C, 14 days) alone or in the presence of a compatible mating partner were analyzed by high performance thin layer chromatography (HPTLC). (**a**–**c**) QM6a and Δ*csg1* asexual (axenic growth) and crossed against CBS999.97 MAT1-1 (CBS1-1); (**d**–**f**) QM6a and Δ*csg2* asexual (axenic growth) and crossed against CBS999.97 MAT1-1 (CBS1-1); Visualization methods of HPTLC results: (**a** and **d**) developed, 366 nm; (**b** and **e**) developed visible light; (**c** and **f**) developed visible light, saturation decreased for better visibility of band patterns. Results shown are consistent in three biological replicates of three pooled plates each. Arrows show regions of altered intensity or presence of bands in Δ*csg2* in the presence of CBS999.97 MAT1-1 (CBS1-1) as compatible mating partner. (**g**) Sexual development in the presence of soybean seedlings**.** CBS999.97 MAT1-1 (CBS1-1) and CBS999.97 MAT1-2 (CBS1-2) as well as mutants of CBS999.97 MAT1-1 lacking the pheromone receptors *hpr1*, *hpr2* or both were grown under mating conditions (daylight, 22 °C) with or without soybean seedlings until fruiting body formation and ascospore discharge. Fruiting bodies indicated by arrows are shown in enlarged sections of the figure. Note that mutants lacking both pheromone receptors are still able to undergo mating and fruiting body formation with a fully fertile mating partner as they can still provide the pheromone precursor required (2). At least three biological replicates per combination were performed and the experiment was performed twice. **(h)** Wild type CBS999.97 MAT1-1 was crossed with Δ*csg1*, Δ*csg2* and respective parental strains at 22 °C, 1700 lux on 2 % MEX medium. Photos were taken after 14 days. No differences in morphology of the fruiting bodies were visible between wild types and deletion mutants.

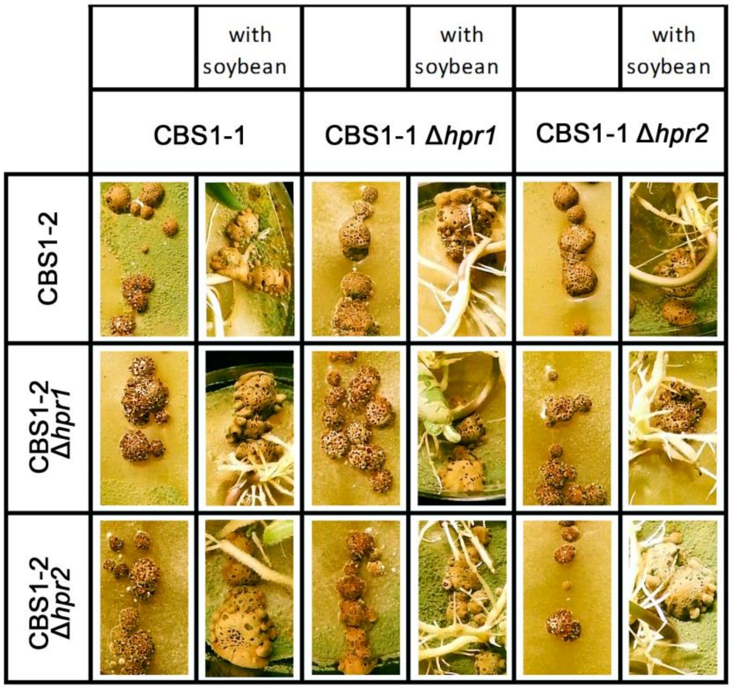


**Figure S6. Fruiting body morphology in the presence of soybean germlings**. CBS999.97 MAT1-1 (CBS1-1) and CBS999.97 MAT1-2 (CBS1-2) as well as mutants in the pheromone receptors *hpr1* or *hpr2* were grown under mating conditions (daylight, 22 °C) with or without soy bean germlings until fruiting body formation and ascospore discharge. Mutants lacking both pheromone receptors are still able to undergo mating and fruiting body formation with a fully fertile mating partner as they can still provide the pheromone precursor required (2). At least three biological replicates were applied per combination and the experiment was repeated twice.

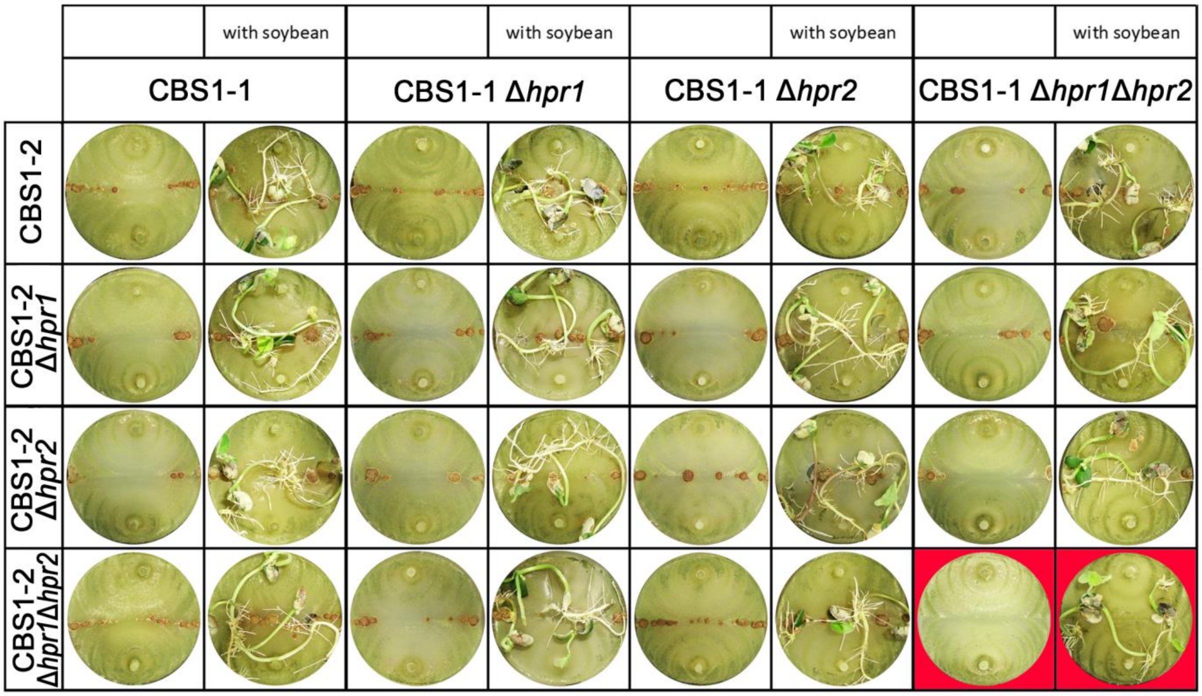


**Figure S7. Fruiting body formation of pheromone receptor mutants in the presence of soy bean germlings.** CBS999.97 MAT1-1 (CBS1-1) and CBS999.97 MAT1-2 (CBS1-2) as well as mutants in the pheromone receptors *hpr1* or *hpr2* or both were grown under mating conditions (daylight, 22 °C) with or without soy bean germlings until fruiting body formation and ascospore discharge. Red background indicates lack of fruiting body formation. Mutants lacking both pheromone receptors are still able to undergo mating and fruiting body formation with a fully fertile mating partner as they can still provide the pheromone precursor required (2). At least three biological replicates were applied per combination and the experiment was repeated twice.

### Supplementary note 2

*Accelerated fruiting body development requires communication with the plant*

Since the presence of plant roots leads to earlier fruiting body formation, we aimed to test whether this is due to communication with the plant or to mere encounter of an obstacle during growth. We found that application of plant root exudate did not cause fruiting body formation to start earlier (Figure S8). In the presence of a toothpick, meant to reflect an altered surface, we found slightly accelerated fruiting body formation, likely in the range of hours (Figure S8). Consequently, accelerated fruiting body formation in the presence of soy germlings is not due to surface alterations or the presence of compounds in the plant exudate, but requires interaction.

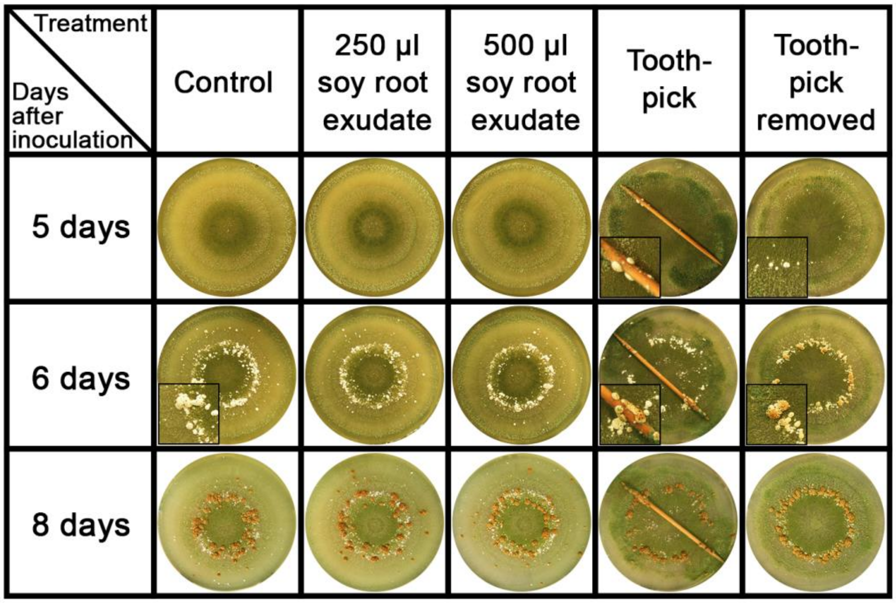


**Figure S8. Sexual development as influenced by plant extracts and toothpicks.** CBS999.97 MAT1-1 and MAT1-2 strains were incubated in light dark cycles at 22 °C on 2 % (w/v) malt extract medium. Same amounts of spore solution of both mating partners were pipetted in the center of the plate to facilitate immediate sensing. Sterile water or soy germling exudate was applied to the center of the plate prior to inoculation, the toothpick was added immediately after inoculation. To evaluate the relevance of diffusible compounds from a toothpick, it was placed on the plate one 24 hours before and then removed prior to inoculation. Fruiting bodies are shown as a close up. One representative picture is shown of five biological replicates.

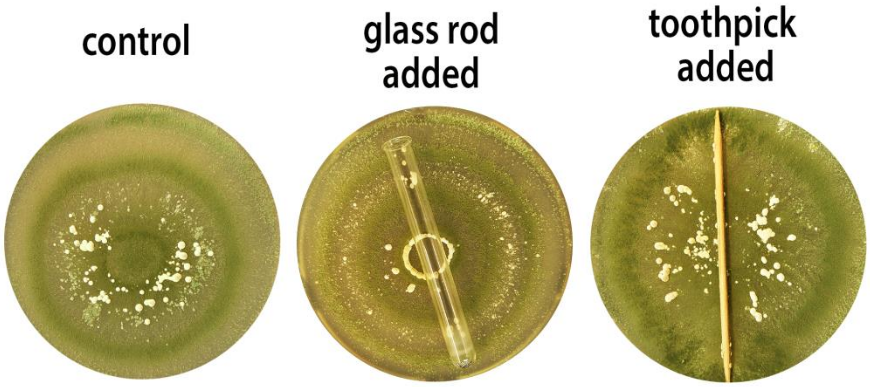


**Figure S9. Sexual development as influenced by an obstacle**. The onset of fruiting body formation is shown. Strains were incubated in light dark cycles at 22 °C on 2 % (w/v) malt extract medium. Same amounts of spore solution of both mating partners were pipetted in the center of the plate to facilitate immediate sensing. The toothpick and the glass rod were added immediately after inoculation. One representative picture is shown of five biological replicates.

**Supplementary note 3**

*Abolished plant sensing does not cause root degradation*

Since plant recognition depends on the presence of pheromone receptors, we reasoned that in their absence, the plant root might only represent a carbon source to be degraded. In this case, a deleterious interaction of the mutants with plant roots in contrast to the wild type could be assumed. We applied the wild type and the double mutants either individually in both mating types or together on the seeds and evaluated the consequences for root and shoot growth in soil cultivations. However, neither root growth nor shoot growth in the presence of the double mutants showed any significant alteration to the wild type treated plants. Only application of both double mutants resulted in a slight decrease in shoot length (Figure S10ab). Consequently, abolished plant sensing does not lead to accelerated degradation of living plant roots or major defects in growth of shoots.

**Supplementary note 4**

*The plant roots for* T. reesei *as a carbon source and sex dev*

The evaluation of the effect of degradable material on the plate (toothpick; Figure S9) also showed, that fruiting bodies formed on the surface of the toothpick, suggesting that the presence of degradable (dead) plant material is relevant. Therefore we repeated the experiment, letting the toothpick soak on the plate prior to application and compared this situation with the presence of glass rods on the surface. Here, fruiting body formation occurred on and around the toothpick, but in the presence of the glass rods only some distribution along the rod was observed that also correlated with some distribution of spores due to capillary forces (Figure S9 toothpick glass). Removal of the toothpick prior to inoculation should show whether extractives from the wood might be involved in fruiting body formation, which was not the case (Figure S8). We conclude that the availability of degradable plant material is relevant for fruiting body formation. This finding is in agreement with the interaction of the fruiting body with the plant root, which shows signs of degradation (Figure 5). Nevertheless, this interaction involving nutrient acquisition by the fungus is not deleterious to the roots.

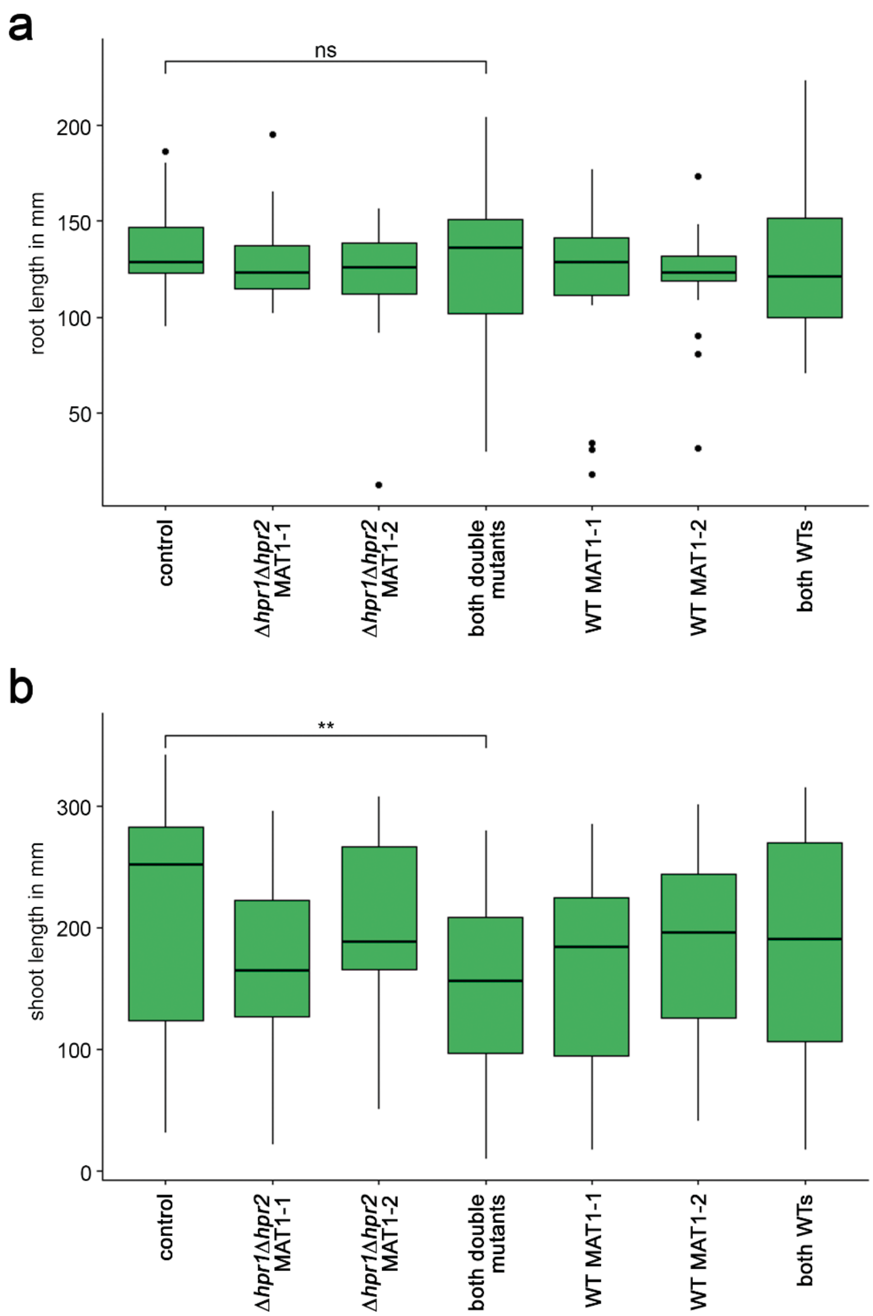


**Figure S10. Impact of *T. reesei* on soybean roots and shoots.** (a) Roots or (b) shoots were measured separately in at least 23 replicates per assay. Length is given in mm. Control = plants alone. Spores of fully-grown petri dishes were applied to the pregerminated seeds by rolling the seeds on the agar surface until full coverage with green conidiospores. Seeds were subsequently transferred to planting pots filled with a mixture of sterile soil, sand and perlite (1:1:1). After 12 days at room temperature, plants were harvested and analyzed.Asterisks mark statistical significance. ** p-value < 0.05.

**Table S1. Strains used in this study**

| strain | mating type | genotype | reference |
|---|---|---|---|
| QM6a | MAT1-2 | wild type | (7) |
| QM9414 | MAT1-2 | wild type | (8) |
| TU-6 | MAT1-2 | wild type | (9) |
| FF1 | MAT1-1 | wild type | (10) |
| FF2 | MAT1-2 | wild type | (10) |
| CBS999.97 MAT1-1 | MAT1-1 | wild type | (3) |
| Δ*hpr1* | MAT1-1 | Δ*hpr1*::*hph*$^{+}$ | (2) |
| Δ*hpr2* | MAT1-1 | Δ*hpr2*::*hph*$^{+}$ | (2) |
| Δ*hpr1*Δ*hpr2* | MAT1-1 | Δ*hpr1*Δ*hpr2*::*hph*$^{+}$ | (2) |
| CBS999.97 MAT1-2 | MAT1-2 | wild type | (3) |
| Δ*hpr1* | MAT1-2 | Δ*hpr1*::*hph*$^{+}$ | (2) |
| Δ*hpr2* | MAT1-2 | Δ*hpr2*::*hph*$^{+}$ | (2) |
| Δ*hpr1*Δ*hpr2* | MAT1-2 | Δ*hpr1*Δ*hpr2*::*hph*$^{+}$ | (2) |
| QM6aΔ*ku80* | MAT1-2 | Δ*ku80*::*amds*$^{+}$ | (11) |
| QM6aΔ*mus53* | MAT1-2 | Δ*mus53*::*bar*$^{+}$ | (12) |
| Δ*cre1* | MAT1-2 | Δ*cre1*::*amdS*$^{+}$ | (13) |
| Δ*gna1* | MAT1-2 | Δ*gna1*Δ*pyr4*::*pyr4*$^{+}$ | (14) |
| Δ*gna2* | MAT1-2 | Δ*gna2*Δ*tku70*Δ*pyr4*::*pyr4*$^{+}$::*hph*$^{+}$ | (15) |
| Δ*gna3* | MAT1-2 | Δ*gna3*Δ*tku70*Δ*pyr4*::*pyr4*$^{+}$::*hph*$^{+}$ | (15) |
| GNA1QL | MAT1-2 | *gna1*$^{QL}$::Δ*pyr4*::*pyr4*$^{+}$ | (14) |
| GNA2QL | MAT1-2 | *gna2*$^{QL}$::*hph*$^{+}$ | this study |
| GNA3QL | MAT1-2 | *gna3*$^{QL}$::Δ*pyr4*::*pyr4*$^{+}$ | (16) |
| Δ*csg1* | MAT1-2 | Δ*csg1*Δ*mus53*::*bar*$^{+}$::*hph*$^{+}$ | (17) |
| Δ*csg2* | MAT1-2 | Δ*csg2*Δ*ku80*::*amdS*$^{+}$::*hph*$^{+}$ | (17) |
| Δ*csg1*Δ*csg2* | MAT1-1 | Δ*csg1*Δ*csg2*::*hph*$^{+}$ | this study |
| c1 | MAT1-1 | wild type, resulted from backcrossing of Δ*csg1* | this study |
| c2 | MAT1-1 | wild type, resulted from backcrossing of Δ*csg2* | this study |
| Δ*acy1* | MAT1-2 | Δ*acy1*::*hph*$^{+}$ | (18) |
| Δ*pkac1* | MAT1-2 | Δ*pkac1*::*hph*$^{+}$ | (18) |
| Δ*gnb1* | MAT1-2 | Δ*gnb1*::*hph*$^{+}$ | (19) |
| Δ*gng1* | MAT1-2 | Δ*gng1*::*hph*$^{+}$ | (19) |
| Δ*phlp1* | MAT1-2 | Δ*phlp1*::*hph*$^{+}$ | (19) |
| Δ4508 | MAT1-2 | Δ4508Δ*ku80*::*amdS*$^{+}$::*hph*$^{+}$ | this study |
| Δ80125 | MAT1-2 | Δ80125Δ*ku80*::*amdS*$^{+}$::*hph*$^{+}$ | this study |
